# A Shifting Landscape: Deconvoluting the Effect of Mg^2+^ and pH on Base Triple Formation in 7SK SL-1

**DOI:** 10.64898/2026.11.19.767663

**Authors:** Lucy Fallon, Harsha Venkatesh, Alisha N Jones

## Abstract

The small-nuclear RNA (snRNA) 7SK is essential for regulating the transcription of genes by RNA polymerase II through its interactions with Hexamethylene Bisacetamide Inducible Protein (HEXIM) and positive transcription elongation factor b (P-TEFb). High-resolution structures of 7SK stem-loop 1 (SL-1) reveal the presence of several base triples that coordinate the interaction with HEXIM. However, different high-resolution structures of 7SK SL-1 exhibit different conformations, each with unique base triple configurations. Previous NMR studies show that the conformation of 7SK SL-1 is sensitive to changes in pH and Mg^2+^ concentration. However, the exact mechanism by which Mg^2+^ and pH influence the formation and stability of base triples in 7SK SL-1 at an atomic level remains unclear. We combine molecular dynamics (MD) simulations, free energy calculations, nuclear magnetic resonance (NMR), and small angle X-ray scattering (SAXS) to conduct a detailed investigation into the mechanism and dynamics of base triple formation in 7SK SL-1, and deconvolute the effects of pH and Mg^2+^. We find that the conformational landscape of 7SK SL-1 dramatically changes depending on the pH and presence of Mg^2+^, and that pH and Mg^2+^ act orthogonally to favor the formation of base triples. We discuss how the dynamic behavior of the base triples may provide a means for 7SK to regulate its interactions with HEXIM, modulating the transcriptional output of RNA polymerase II.

**SIGNIFICANCE STATEMENT:** The small nuclear RNA 7SK stem-loop 1 (SL-1) is a master regulator of RNA transcription by RNA polymerase II, and harbors several base triples with a protonated third base that make up the binding platform for the protein HEXIM. Here, we marry simulation and experiment to quantify the shift in the conformational ensemble of the RNA 7SK SL-1 in response to the protonation state and chemical environment of the base triples through changes in the free energy surface, free of protein partners. Our results present an atomic-detailed framework for how environmental factors can dramatically reshape biomolecular free energy surfaces, and further advance our textbook knowledge of the role of alternative protonation states in RNA structure and dynamics.

## INTRODUCTION

The small nuclear RNA (snRNA) 7SK is a highly abundant and evolutionarily conserved RNA that acts as a transcriptional regulator for RNA polymerase II (Pol II) (1–3). 7SK tightly regulates the activity of the positive transcription elongation factor b (P-TEFb) by sequestering it in a ribonucleoprotein (RNP) complex through interactions with Hexamethylene Bisacetamide Inducible Protein (HEXIM) (4–12). P-TEFb binds HEXIM, while a HEXIM-forming dimer binds to stem-loop 1 (SL-1) of 7SK (13). Release of P-TEFb from the HEXIM-7SK RNP complex allows P-TEFb to rescue Pol II from its stalled state, continuing transcription (14). The Human Immunodeficiency Virus (HIV) trans activator (Tat) protein outcompetes HEXIM and binds 7SK SL-1 to hijack PTEF-b for transcription of its viral transcriptome (15, 16). Structural studies have revealed the presence of multiple base triples in 7SK SL-1 arranged into arginine sandwich motifs (ASMs), which are recognized and engaged by the arginine-rich motifs (ARM) of either Tat or HEXIM (12, 16, 17).

Multiple high-resolution structures of 7SK SL-1, with variable apical loops, have been reported (12, 16–18). Interestingly, the base triple configurations differ across the experimental structures. One NMR study (PDB ID 6MCI) reported 7SK SL-1 in a compact state, harboring four base triples arranged into three ASMs, and a pseudo-ASM (pASM; **Figure 1A**) (16). This base triple configuration is present in structures of 7SK SL-1 bound to the ARMs of Tat or HEXIM, suggesting it may be a binding-competent conformation necessary for recognition and binding of the Tat and HEXIM ARMs (12, 16). The structure of 7SK SL-1 has also been determined by X-ray crystallography: compact states, harboring either six or seven base triples have been resolved (PDB IDs 5LYU, 5LYS) (17). Finally, another NMR structure (PDB ID 5IEM) shows 7SK SL-1 can adopt an extended conformation, where no base triples are formed (18).

**Figure 1.**
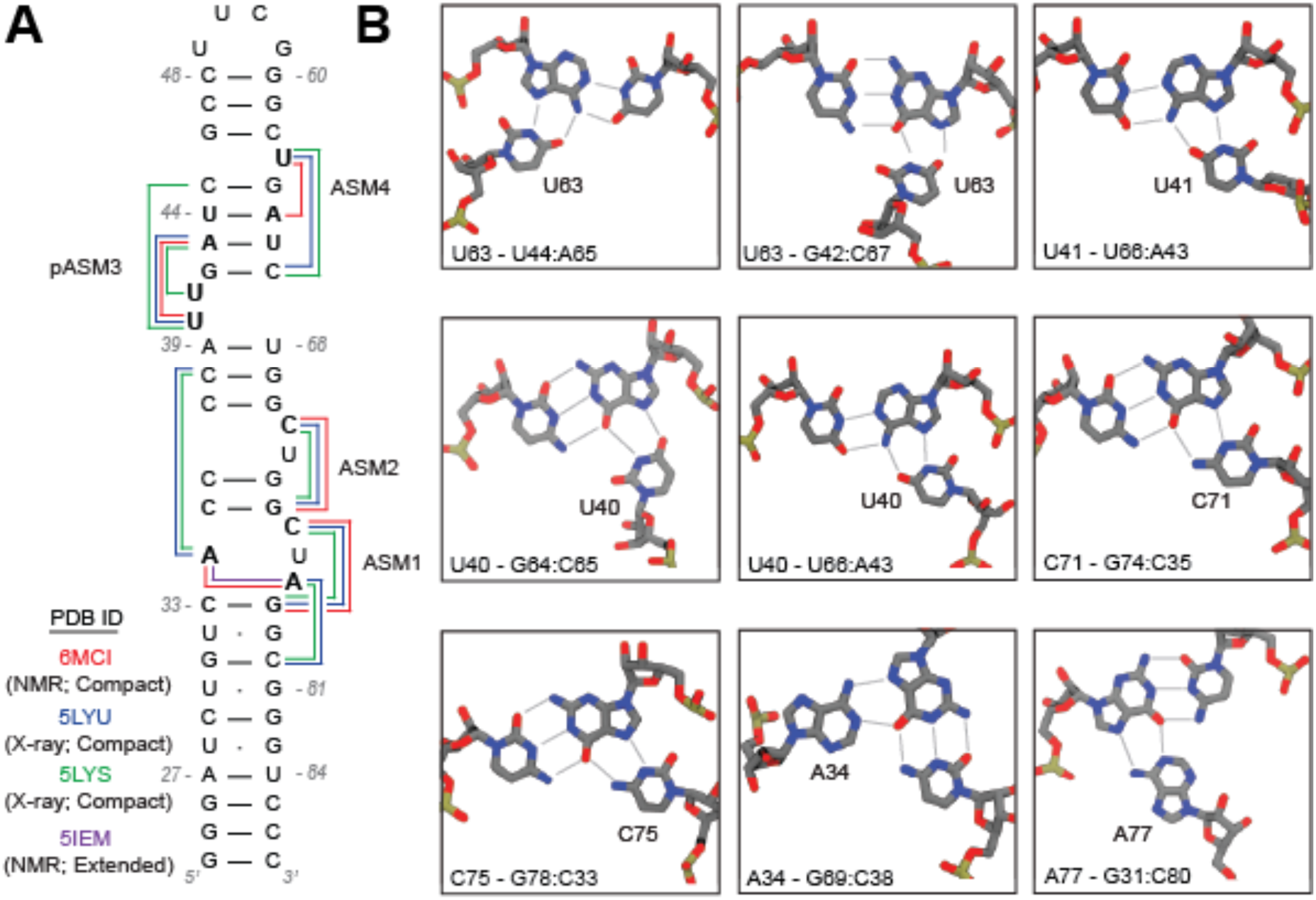
**A** Secondary structure of 7SK SL-1 with the tertiary features across the various high-resolution structures represented, along with their respective PDB IDs. Base triples and non-canonical interactions are indicated as colored lines, colored according to the experiment. **B** Visualizations of the base triples seen in the different high-resolution structures of 7SK SL-1.

Several reports have proposed that the third base (bold) of the **A** - G:C and **C** - G:C base triples in the aforementioned structures are protonated (A34, C71, C75, and A77) (17, 19, 20). Protonation of C71 and C75 was experimentally supported in the compact NMR-resolved structure (PDB ID 6MCI) by the characteristic downfield shift of the N4 amino protons (16). The other structures of 7SK SL-1 in the compact state also harbor the C71 and C75 base triples; the N3 atoms of these nucleotides are less than 3 Å away from the N7 atoms of the Watson-Crick-Franklin base-paired guanines, a distance that is sufficient to support a protonated state (12, 17, 19).

The two A - G:C base triples are only observed in the crystal structures; the N1 atom of the adenines are positioned 2.6 Å away from the O6 atom of the proximal guanines, introducing an unfavorable steric clash that would potentially be relieved by the presence of a proton (17). In the HEXIM-bound structure, A34 and A77 are not involved in base triples but are instead hydrogen bonding with each other in an A-A mismatch, forming the interior of the first ASM (ASM1). This A-A mismatch, but not the ASM, is further observed in the extended conformation.

Each of the high-resolution structures were resolved in different buffer conditions, which is the source of the differences in tertiary structure (17, 19). Available biophysical data suggests that the conformational landscape of 7SK SL-1 is dependent on pH and the concentration of Mg^2+^ (17, 19). Two different NMR studies observed chemical shift perturbations (CSPs) of several resonances after the addition of Mg^2+^, including that of the C71 - G74:C35 base triple (18, 19). The 2016 NMR study reported that the U41 - A43:U66 base triple only formed in their high (3 mM) MgCl_2_ model (18). An X-ray structure of 7SK SL-1 resolved a hexahydrated Mg^2+^ ion near the base triples involving C71 and A34, potentially stabilizing the two base triples, while Small-Angle X-ray Scattering (SAXS) data demonstrated that Mg^2+^ drives 7SK SL-1 into a more compact conformation (17, 19).

It has been proposed that the shift from the extended state to the compact state is likely driven by protonation of C71 and C75, and the subsequent formation of base triples – which are further stabilized by Mg^2+^ (19). The NMR structure solved by Pham *et al.* showed the binding-competent conformation as the dominant structure, whereas a molecular dynamics (MD) simulation-focused study suggested that 7SK SL-1 spontaneously converts between the different experimental structures as a means to regulate interactions with HEXIM (16, 20). However, this simulation study was performed on a shortened model of 7SK SL-1 lacking the C - G:C and A - G:C base triples, preventing any assessment of their relative stabilities and sensitivities towards buffer conditions (20). To this end, a study that couples pH, MgCl_2_, and the full length structured RNA is necessary to cull these disparate views.

In this study, we combine MD simulations, free energy calculations, NMR, and SAXS experiments to probe the mechanism of base triple formation in 7SK SL-1, and characterize the influence that Mg^2+^ and pH have on the conformational landscape of 7SK SL-1. We find that Mg^2+^ and pH have orthogonal, synergistic effects on 7SK SL-1: Mg^2+^ modulates the global conformation of 7SK SL-1, shifting it towards a more compact conformation at increased concentrations, and orienting the RNA backbone in conformations amenable for base triple formation. Reducing pH has little effect on the global compaction of 7SK SL-1, or the stability of the U40, U41, or U63 base triples – but it does drive formation of the C71 and C75 base triples, and subsequently the ASMs. Our results help rationalize experimental observations on the role of Mg^2+^ and pH on the dynamics of 7SK SL-1, and provide a mechanism for base triple formation in 7SK SL-1. We discuss the implications of our results on the binding of HEXIM and regulation of P-TEFb.

## RESULTS

### MD Simulations Suggest the Stability of the Base Triples are Sensitive to pH & Mg^2+^

As an initial assessment on the sensitivity of the base triples to pH, we first performed triplicate 2 µs MD simulations of 7SK SL-1 (PDB ID 5LYU) at both low and neutral pH in the presence and absence of Mg^2+^. The “low pH” simulations described in this text refer to 7SK SL-1 with A34, C71, C75, and A77 in the protonated state (referred to in this text as the “ionizable base triples”). Our simulations at “neutral pH” correspond to simulations with A34, C71, C75, and A77 in the neutral state, lacking the extra proton.

We monitored the distance between the third base and the Hoogsteen face of the participating nucleotide in the base triples throughout our simulations in the presence of Mg^2+^ (**Figure 2**). In agreement with previous NMR studies, the four ionizable base triples in 7SK SL-1 are more stable in our simulations at low pH, when the third base (C or A) harbors the extra proton (**Figure 2C-F**) (16, 19). This is indicated by the large peaks centered around 2.5 Å, which shift to larger distances at neutral pH. When simulating this system at neutral pH, the base triple conformation is abandoned and is not sampled throughout the rest of the simulations (**Figure 2C-F**). The two non-ionizable base triples (U40 - A43:U66 and U63 - G42:C67) in this system do not appear sensitive to the protonation state of C71, C75, A34, and A77, and behave nearly identically in our simulations at both low and neutral pH (**Figure 2A, B**).

**Figure 2.**
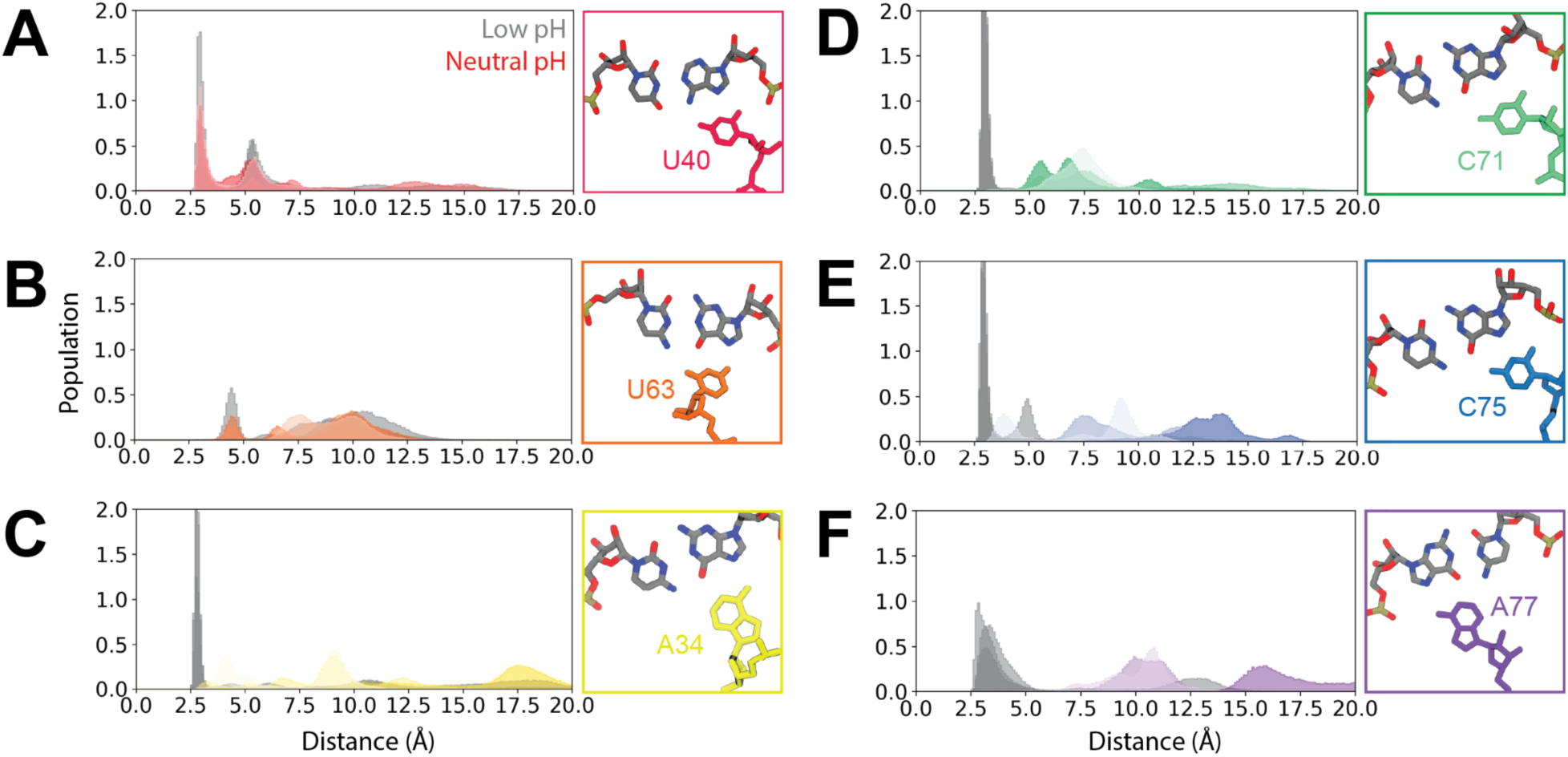
Distance distributions of the base triples throughout 2.0 µs MD simulation of 7SK SL-1 starting from the X-ray structure (PDB ID 5LYU), at both low (grey) and neutral (colored) pH values (17). The colors of the distance plots match the color scheme of the base triples from Figure 1B. The plots show the distance distribution of **A** U40 - A43:U66, **B** U63 - G42:C67, **C** A34 - C38:G69, **D** C71 - G74:C35, **E** C75 - G78:C33, and **F** A77 - G31:C80. The structure of each base triple is shown inset to the right of each distance plot, taken from the X-ray structure.

The same behavior was seen for our simulations of the other two compact structures. The non-ionizable base triples are not sensitive to pH (**Supplemental Figure 1**), whereas the four ionizable base triples are significantly less populated at neutral pH than when protonated (**Supplemental Figure 1**). The extended NMR structure of 7SK SL-1 does not adopt any of the base triples throughout the simulation (**Supplemental Figure 1**), likely because the length of our simulations (2.0 µs) is shorter than the exchange rate of 7SK SL-1 alternating between the extended and compact states. Our initial MD results suggest that pH has a significant effect on the stability of the ionizable base triples across each of the three compact structures, but not on the base triples involving U40, U41, or U63.

Conversely, the absence of Mg^2+^ destabilizes the non-ionizable base triples and does not significantly affect the stability of the four ionizable base triples in our simulations. The non-ionizable base triples (involving U40, U41, and U63) exhibit a dramatically broadened distance distribution at both low and neutral pH values in the absence of Mg^2+^ (**Supplemental Figure 2**). The ionizable base triples maintain the native base triple state for the duration of the simulations at low pH, both in the presence and absence of Mg^2+^ (**Supplemental Figure 2**). The increased stability of the non-ionizable base triples in the presence of Mg^2+^ in our simulations is consistent with previous studies on 7SK SL-1 – and with NMR experiments of DNA base triples, which have shown that ionic strength has more of a stabilizing effect on base triples where the third base is a T (U), compared to base triples where the third base is a C or A (18, 21). The effect of pH or Mg^2+^ on the stability of the base triples depends on the exact identity of the nucleotides involved.

### Mg^2+^ Ions Stabilize the Compact State of 7SK SL-1

Previous work has shown the presence of Mg^2+^ can help drive 7SK SL-1 towards the compact state (19). We monitored the radius of gyration (*R_G_*) of 7SK SL-1 in our MD simulations to understand the effect that Mg^2+^ has on the compaction of 7SK SL-1.

Changing pH has very little effect on the *R_G_* in our simulations. Regardless of the presence of Mg^2+^, shifting from low pH to neutral pH does not change the *R_G_* for each of the four systems: the *R_G_* is centered around 20-22 Å for compact systems (PDB IDs 5LYU, 5LYS, 6MCI), and 18-26 Å in the extended state (PDB ID 5IEM; **Figure 3**).

**Figure 3.**
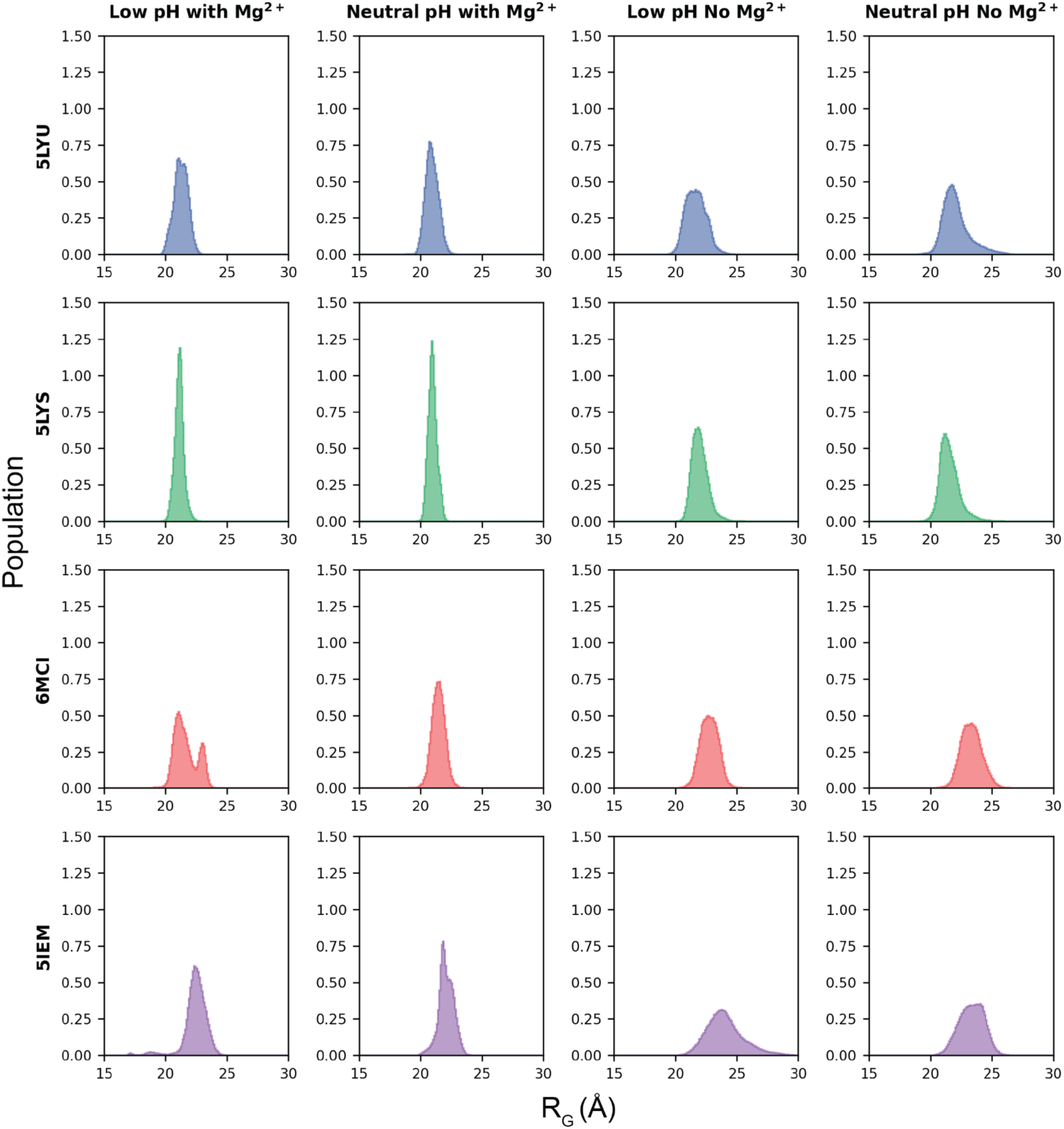
Radius of gyration of 7SK SL-1 in our triplicate 2.0 µs MD simulations starting from the compact X-ray structures (PDB IDs 5LYU, 5LYS), the compact NMR structure (PDB ID 6MCI), or the extended NMR structure (PDB ID 5IEM), at low and neutral pH in the presence and absence of Mg^2+^.

The presence of Mg^2+^ ions, however, has a strong effect on the global compaction of 7SK SL-1. We observe a distinct shift in the *R_G_* when comparing our simulations with and without Mg^2+^. At low pH with Mg^2+^ the *R_G_* is centered around 20-22 Å for the compact systems (PDB IDs 5LYU, 5LYS, 6MCI), and 18-24 Å for the extended system (PDB ID 5IEM; **Figure 3**). But at low pH without Mg^2+^ the *R_G_* distribution broadens to 20-24 Å for the compact systems, and 20-26 Å for the extended system. At neutral pH the effect of Mg^2+^ on the compaction of 7SK SL-1 is even more apparent: in the presence of Mg^2+^ the *R_G_* distribution is centered around 20-22 Å for the compact systems, and 22-24 Å for the extended system, but is broadened to 20-26 Å for the compact systems, and 21-30 Å for the extended system in the absence of Mg^2+^.

To gain a structural understanding of how Mg^2+^ drives 7SK SL-1 towards a compact state, we searched for regions of high Mg^2+^-occupancy in our simulations. We identified four regions consistently occupied by Mg^2+^ ions that appear to stabilize the compact state (**Figure 4A**).

**Figure 4.**
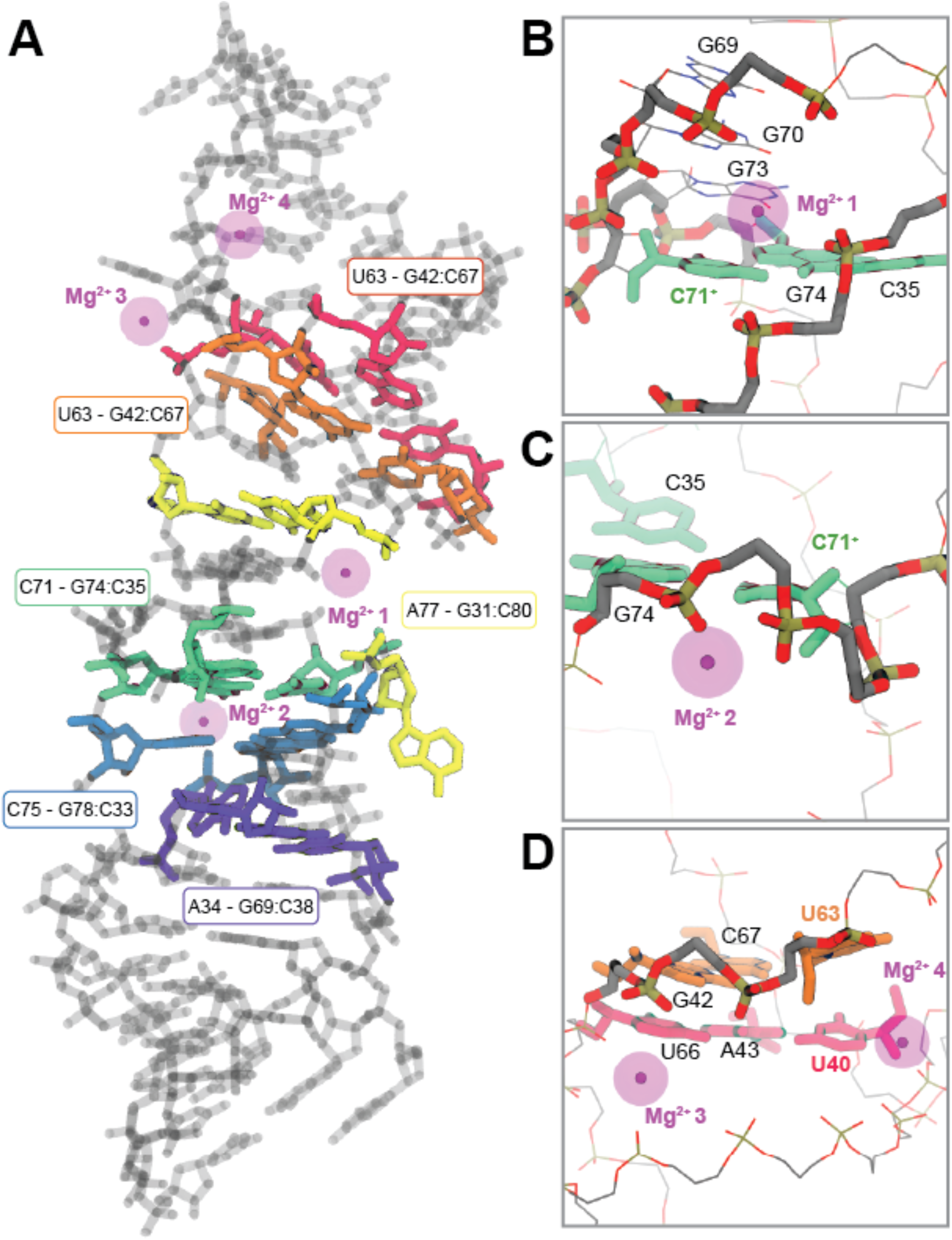
Regions of high Mg^2+^ occupancy in our MD simulations of 7SK SL-1 at low pH in the presence of Mg^2+^. **A** Snapshot of a simulation structure from our simulations of 7SK SL-1 starting from the X-ray structure (PDB ID 5LYU), with base triples colored according to Figure 1, and Mg^2+^ ions in magenta. **B** A high-occupancy Mg^2+^ ion that interacts with the phosphate backbone across the major groove through its hydration sphere, crimping the RNA strands together. **C** A second high-occupancy Mg^2+^ ion that coordinates to the backbone of the RNA, and stabilizes the kink in the RNA backbone near the C71 - G74:C35 base triple. **D** Two high-occupancy Mg^2+^ ions near the base triples involving U40 and U63, that interact with the phosphate backbone across the major groove through their hydration spheres, crimping the RNA strands together, and compacting the RNA.

The first region of high Mg^2+^ occupancy seen in our simulations lies in the same position as the Mg^2+^ ion resolved in the crystal structure, adjacent to the base triples involving C71^+^ and A34^+^ (17). This Mg^2+^ ion does not coordinate directly to the RNA, and instead forms interactions through its hydration sphere, bridging the phosphate backbone across the RNA major groove (**Figure 4B**) (17). The Mg^2+^ ion at this position exchanges rapidly in our simulations, yet we observe near constant occupancy at this site. This site is proximal to four sequentially stacked guanines (G69, G70, G73, and G74; **Figure 4B**), the Hoogsteen faces of which create a partially negatively charged channel that appears to attract the Mg^2+^ ion to this region. Notably, three of these four guanine nucleotides (G69, G70, and G73) exhibited CSPs in response to the addition of Mg^2+^ in a previous NMR study; G74 does not, possibly because its Hoogsteen face is occluded by C71^+^ when engaged in the base triple (18, 19). As observed in the original X-ray study, the Mg^2+^ ion at this position appears to rigidify the central region of 7SK SL-1 by linking the RNA backbone across the major groove through its hydration sphere (17).

The second high-occupancy Mg^2+^ site is near the C71^+^ - G74:C35 base triple. The Mg^2+^ ion at this site coordinates directly to the oxygen atom of the phosphate backbone of G74 in our simulations, and compacts the RNA by stabilizing the kink in the RNA backbone spanning nucleotides C71 to G74 (**Figure 4C**). This Mg^2+^ ion helps to alleviate the electrostatic strain from the proximal phosphate groups resulting from the kinked backbone. Compaction of the RNA backbone then orients C71 in a position where it is able to form a base triple with the G74:C35 base pair.

Lastly, we observe two regions of high Mg^2+^ occupancy near the non-ionizable base triples involving U40, U41, and U63 (**Figure 4D**). The Mg^2+^ ions at these two sites rapidly exchange, but consistently localize in the same regions throughout the course of the simulations. These two Mg^2+^ ions interact with the RNA through their hydration spheres, and rigidify the upper portion of 7SK SL-1. In the simulations starting from the X-ray structures, one of these two Mg^2+^ ions stabilizes the kink in the RNA backbone at U63, which helps constrain it in a position amenable for base triple formation with the G42:C67 base pair (left ion in **Figure 4D**). The other Mg^2+^ ion, near U40, provides a bridging interaction across the RNA major groove through coordinating water molecules, which compacts the upper region of the RNA by crimping the RNA backbone together (right ion in **Figure 4D**).

Altogether, our simulation data suggests that Mg^2+^, not pH, drives 7SK SL-1 towards a more compact state, and stabilizes the U40, U41, and U63 base triples – congruent with observations from previous NMR, SAXS, and UV-melting experiments (18, 19).

### The pKa Values of the Third Bases Are Elevated for the Ionizable Base Triples

Protonation of the third bases increases the stability of the ionizable base triples in our MD simulations. However, 7SK exists in the nucleus, which is near neutral pH, and the *pKa* of free cytosine is far from neutral at 4.2; free adenosine is even lower at 3.6 (22). It is therefore unclear whether these nucleotides would be protonated inside the cell at physiological pH, and whether the ionizable base triples would be present in significant populations. Past NMR studies on complex RNA structures have confirmed the presence of protonated cytosines, with apparent *pKa* shifts well above neutral, sometimes even as high as 8.1 (23–27). We hypothesized that the *pKa* of the third base in the four ionizable base triples must be elevated towards neutral to facilitate stable formation of the base triples at physiological pH. We estimated the *pKa* change of the third base in each of the four ionizable base triples in 7SK SL-1 using thermodynamic integration (TI) (**Supplemental Figure 3**) (28). This methodology has been successfully applied in the past for the calculation of *pKa* changes in nucleic acids and amino acids (29–32).

In the compact structure of 7SK SL-1 (PDB ID 5LYU), the *pKa* of C71 and C75 are 6.3 and 7.3, respectively (**Figure 5**): 2 - 3 pH units above the value of 4.2 for free cytosine (22). The two adenine nucleotides also show significantly elevated *pKa* values: 6.2 for A34, and 5.5 for A77: approximately 2 - 2.5 pH units above the reference value of 3.6 for free adenosine (22). This suggests that the C - G:C base triples are populated at physiological conditions, whereas the A - G:C base triples are likely rare excited states due to the *pKa* of A34 and A77 being much further from neutral.

**Figure 5.**
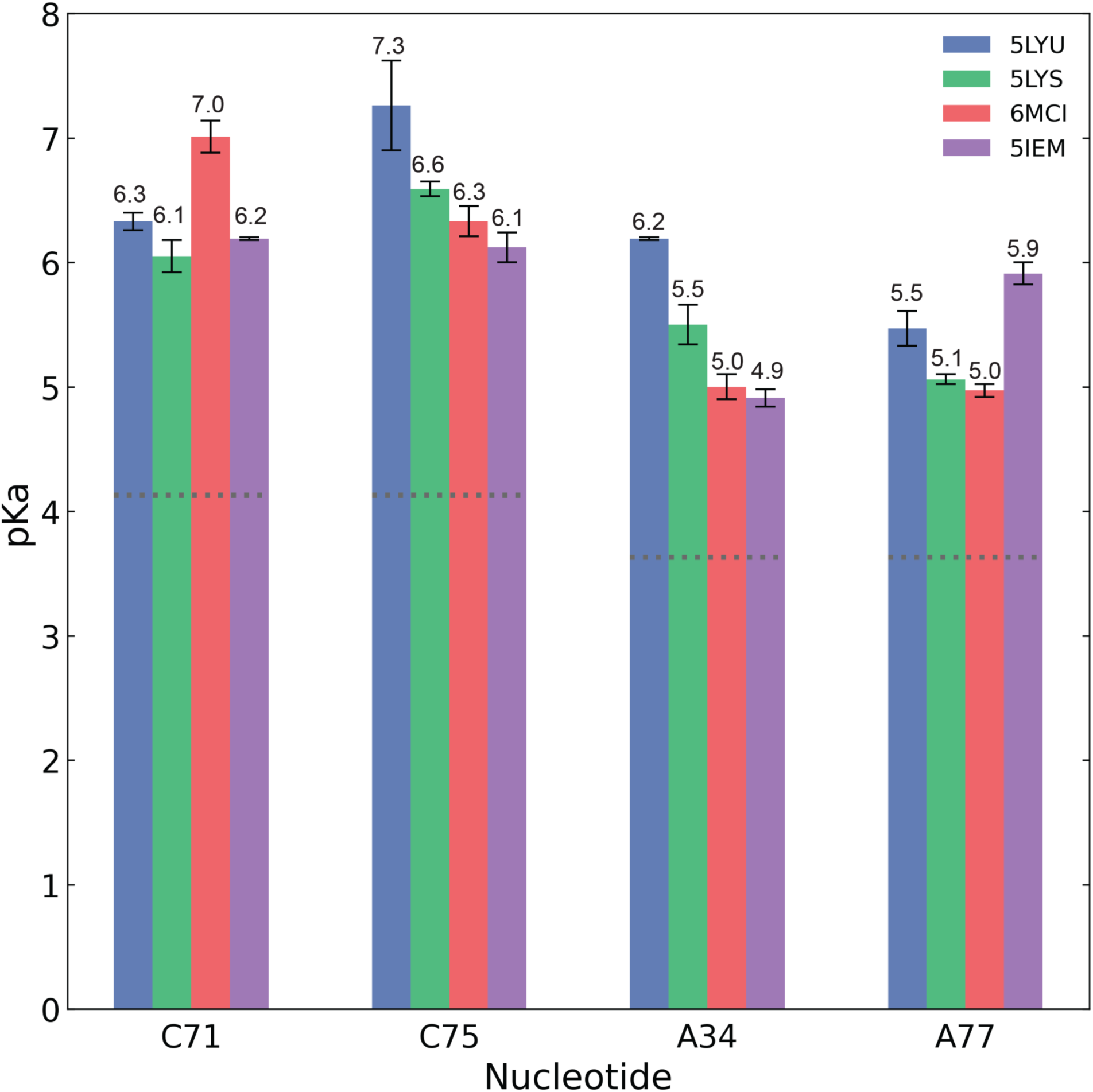
The calculated *pKa* values for C71, C75, A34, and A77 in the four ionizable base triples from two 50 ns replicates, with standard deviations. The *pKa* values were calculated for the third bases starting from each of the four experimental structures. The dotted gray lines represent the reference *pKa* values for the free nucleotide.

We also calculated the *pKa* of A34, C71, C75, and A77 in each of the remaining three experimental structures, given that this parameter is highly dependent on its chemical environment (17). We found that the *pKa* of each nucleotide is within ∼1.0 *pKa* unit relative to 5LYU (**Figure 5**). For example, C71 (*pKa* 6.3) shows a *pKa* value of 6.1 when the calculation was done using another X-ray structure (PDB ID 5LYS), but is elevated to 7.0 when using the compact solution NMR structure (PDB ID 6MCI) (16, 17).

Altogether, our *pKa* calculations strongly suggest that the *pKa* of the third base in each of the four ionizable base triples is significantly elevated towards neutral. The nucleotides A34, C71, C75, and A77 exhibit *pKa* shifts of up to 6.2, 7.0, 7.3, and 5.9, respectively, resulting in a *pKa* shift of nearly 2 - 3 units above the pKa values of the free nucleotide. According to these values, at the near-neutral pH inside the cell nucleus, C71 and C75 could spontaneously exist in the protonated state, favoring formation of the C - G:C base triples. Whereas A34 and A77 likely exist predominantly in the neutral state, since the calculated *pKa* values do not reach neutral. The A34 and A77 base triples may be alternative excited states that are less frequently populated by 7SK SL-1, which would explain why they are not observed in any of the solution NMR structures, or when Tat / HEXIM are bound (12, 16–18).

### The Free Energy Surface of 7SK SL-1 Changes Dramatically Depending on pH and Mg^2+^

To quantify the extent to which pH and Mg^2+^ influence the stability of the base triples and the global conformation of 7SK SL-1, we used Well-Tempered Adaptively Biased MD (WT-ABMD) to map the free energy surface of base triple formation in 7SK SL-1 (**Supplemental Note 2**) (33–36). We used two collective variables (CVs) to calculate the potential of mean force (PMF) of base triple formation: (i) the coordination number (*CN*), to track base triple formation, and (ii) the *R_G_*, as a measure of overall compactness. We chose these CVs because our MD simulations confirmed that these two observables are sensitive to pH and Mg^2+^ concentration.

At low pH with Mg^2+^, the free energy surface of 7SK SL-1 harbors three global minima (**Figure 6A**): the two X-ray structures (PDB ID 5LYU & 5LYS), and a compact structure with no base triples at *CN=0*, *R_G_=22-23 Å*. Under these conditions, 7SK SL-1 strongly prefers the compact state: all three minimum energy structures lie in compact regions of *R_G_* (21-23 Å). The two X-ray structures are equivalent in energy and separated by a 2.5 kcal/mol energy barrier, suggesting the RNA may spontaneously alternate between these two conformations. The extended (PDB ID 5IEM) and compact (PDB ID 6MCI) NMR structures are energetically equivalent, lying ∼10.0 kcal/mol above the minimum energy structures. The ionizable base triples are strongly populated, with the A34, C71, and C75 base triples being sampled the most, whereas the A77 base triple is less populated (**Supplemental Figure 4A**). The non-ionizable base triples are more dynamic, but the U40 - A43:U66 base triple is most populated, followed by the U41 - A43:U66 and U40 - G64:C45 configurations. The two base triples involving U63 are sampled the least, suggesting they are the least stable of the different base triples observed in 7SK SL-1.

**Figure 6.**
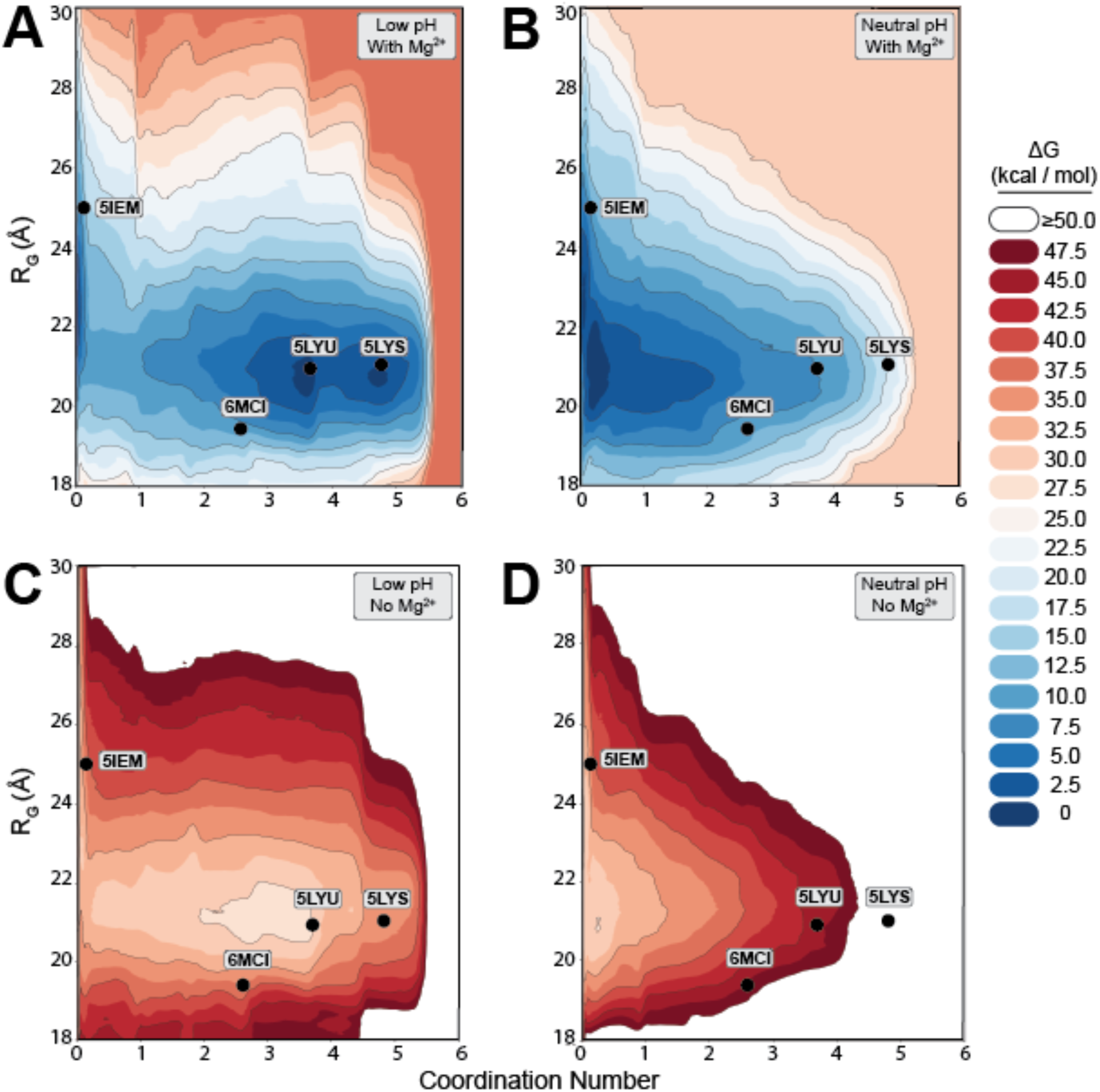
PMFs of 7SK SL-1 alternating between the four different experimental structures with Radius of Gyration (R_G_) and Coordination Number as the two CVs at **A** low pH with Mg^2+^, **B** neutral pH with Mg^2+^, **C** low pH without Mg^2+^, and **D** neutral pH without Mg^2+^. The black circles indicate the values of the experimental reference structures, and their PDB IDs are labeled for clarity. Each color change represents a 2.5 kcal/mol change in free energy, while contour lines represent a 5.0 kcal/mol change in energy.

Next, we estimated the effect that shifting to neutral pH has on the conformational landscape of 7SK SL-1. At neutral pH in the presence of Mg^2+^ the free energy surface collapses (**Figure 6B**), and is characterized by two energetic minima: one corresponding to an intermediately-compact structure with no base triples (*CN=0, R_G_=22-23 Å*), and a second larger minimum-energy well corresponding to a compact structure with one non-ionizable base triple (*CN=0.1-0.5*, *R_G_=20-22 Å*). In these conditions, the ionizable base triples are not populated at all (**Supplemental Figure 4B**), each showing distances greater than 5.0 Å. The non-ionizable base triples behave nearly identically as in the simulations at low pH: the U40 - A43:U66 base triple is most populated, followed by the U40 - G64:C45 and U41 - A43:U66 base triples.

To dissect the effect that Mg^2+^ has on the conformational landscape of 7SK SL-1, we next calculated the free energy surface *without* Mg^2+^ – again at both low and neutral pH values. At low pH without Mg^2+^ (**Figure 6C**) the free energy surface is qualitatively similar to that with Mg^2+^ (**Figure 6A**): the X-ray structure (PDB ID 5LYU) sits at the global minimum, but is no longer energetically degenerate with the other crystal structure (PDB ID 5LYS), and is instead ∼7.5 kcal/mol away from it. The compact and extended NMR structures are still energetically equivalent (PDB IDs 6MCI & 5IEM), and are ∼10.0 kcal/mol higher in energy than the minimum energy structure – as in our free energy surface at low pH with Mg^2+^ (**Figure 6A**). There are no significant differences in the populations of the ionizable base triples at low pH upon removing Mg^2+^ (top four plots of **Supplemental Figure 4A**, **C**). However, the non-ionizable base triples all show a dramatic decrease in their populations in the absence of Mg^2+^ (orange, red, black, magenta plots in **Supplemental Figure 4A, C**). Additionally, the *R_G_* distribution notably broadens upon removal of Mg^2+^ at low pH (last plot, light blue, in **Supplemental Figure 4A, C**).

At neutral pH in the absence of Mg^2+^ the free energy surface collapses as before, and the minimum energy basin lies at *CN=0* between *R_G_=20-25 Å*, corresponding to structures with no base triples, ranging from compact to extended (**Figure 6D**). In these conditions 7SK SL-1 appears able to alternate between the extended and compact states with no energetic penalty, based on the flat *R_G_* at *CN=0-0.5*. However, the formation of base triples becomes increasingly unfavorable as the *CN* CV increases. The ionizable base triples are not sampled at all at neutral pH, and the *R_G_* and distance distribution of the non-ionizable base triples show the same broadening behavior as at low pH without Mg^2+^ (**Supplemental Figure 4D**): regardless of pH, the absence of Mg^2+^ broadens the *R_G_* and decreases the populations of the non-ionizable base triples.

Since our TI calculations estimate the pKa of both C71 and C75 near ∼7.0, and A34 and A77 near ∼6.0, we asked which base triple configuration is most favorable if only C71 and C75 were protonated, to mimic the near-neutral pH found inside the cell nucleus. We calculated the free energy surface of base triple formation again with only C71 and C75 protonated. Under these conditions, the free energy surface changes dramatically (**Supplemental Figure 5**): the binding- competent base triple configuration is lower in energy than the X-ray structures, and an intermediate-compact state with no base triples sits at the global minimum. Under these conditions, 7SK SL-1 can potentially spontaneously alternate between an extended state with no base triples, to the binding-competent conformation with base triples.

### Formation of the Base Triples Requires Compaction of the RNA Backbone

We next assessed whether there are local backbone conformations specific to base triple formation. In all the experimental structures of 7SK SL-1, the third base in each of the base triples lies at a bulged nucleotide possessing a kink in the RNA backbone, that appears to position the third base for hydrogen bonding (12, 17, 19). We hypothesized that formation of the base triples restricts the RNA backbone local to the third nucleotide in specific conformations. To test our hypothesis, we monitored the pseudo-dihedral angles eta (***η***) and theta (***Θ***) throughout our ABMD simulations (37). These pseudo-dihedral angles simplify the six dihedrals of the RNA backbone into two pseudo-torsions, allowing for the visualization of “Ramachadran-like” plots.

We separated our ABMD ensemble into two pools: one in which the C71 - G74:C35 base triple is present, and another where it is absent, and calculated the ***η* / *Θ*** distribution across these two pools for C71 and the two nucleotides immediately following it. We found that when C71 is engaged in the base triple the ***η* / *Θ*** distributions for C71, U72, and G73 are constrained to certain values (**Figure 7A**). When C71 is not participating in a base triple, the ***η* / *Θ*** distributions expand, and these three nucleotides sample a broad range of backbone conformations (**Figure 7B)**.

**Figure 7.**
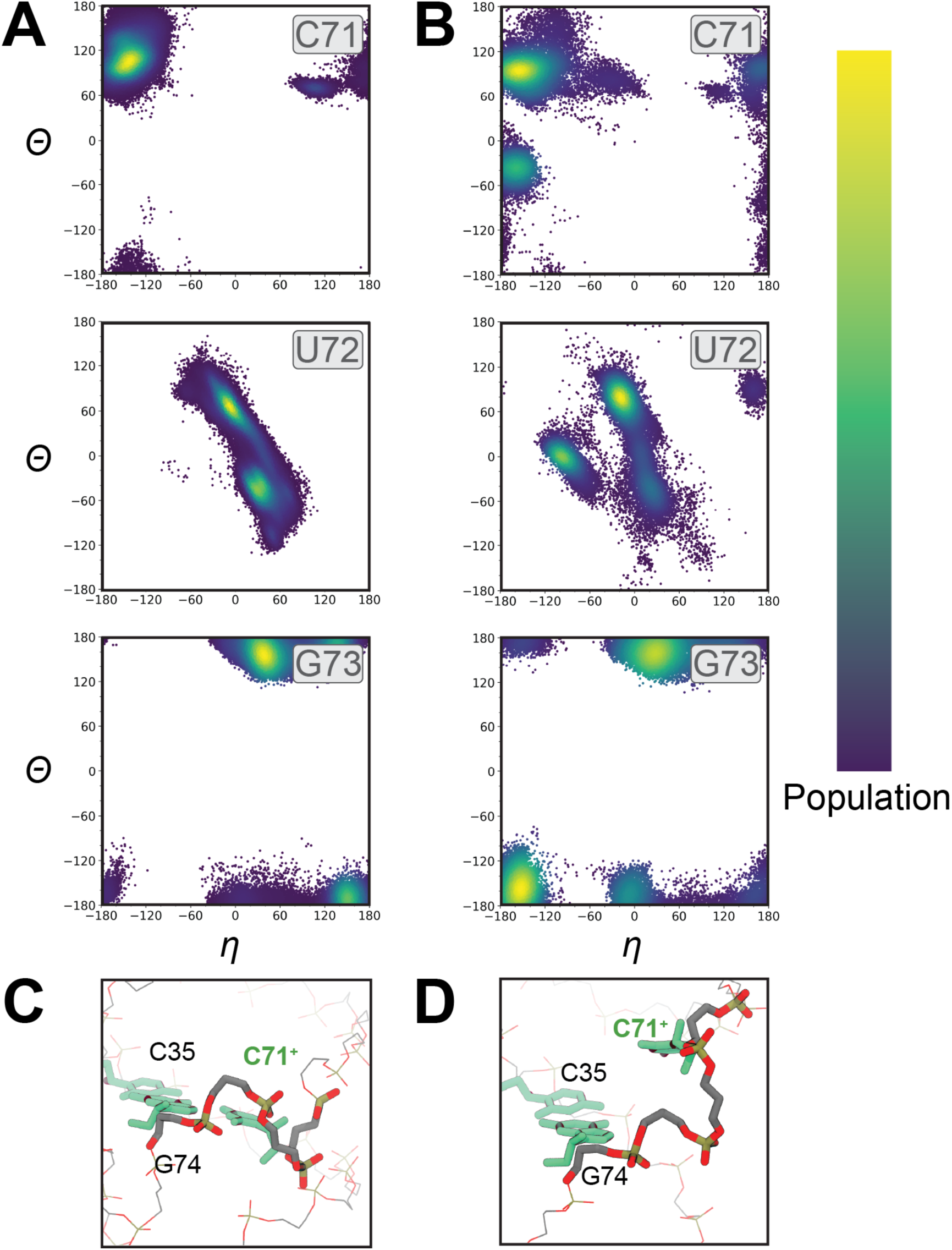
Eta / theta (***η* / *Θ***) distributions of C71, U72, and G73 for structures from our WT-ABMD simulations in which the C71 base triple is **A** present, or **B** absent. The population of each value is colored according to intensity. **C** The backbone conformation of C71, U72, and G73 as seen in the compact NMR structure (PDB ID 5LYU) when C71 is in the base triple conformation. **D** The backbone conformation of C71, U72, and G73 as seen in the extended NMR structure (PDB ID 5IEM) when C71 is not in the base triple conformation.

We observed the same trend for the base triples involving C75, A34, and A77: structures where a base triple is present exhibit a significantly narrower ***η* / *Θ*** distribution than structures without the base triple (**Supplemental Figures 6-8**). Base triple formation in 7SK SL-1 is only permitted from certain backbone configurations of the nucleotides local to the third base. The exact conformation of the RNA backbone needed for the base triple to form is not universal and depends on the position of the third base relative to its participating base pair in sequence-space, as reflected by the different ***η* / *Θ*** distributions. For each of the base triples in 7SK SL-1, rearrangement of the RNA backbone is required to position the third base for hydrogen bonding with its participating base pair.

### SAXS Measurements Show a Compaction of the RNA at Low pH in the Presence of Mg^2+^

We used SAXS to dissect the effects of Mg^2+^ and pH on the global conformation of 7SK SL-1. We measured SAXS scattering profiles under four buffer conditions: (i) low pH with Mg^2+^ (pH 4.5, 4 mM MgCl_2_), (ii) neutral pH with Mg^2+^ (pH 7.0, 4 mM MgCl_2_), (iii) low pH without Mg^2+^ (pH 4.5, 0 mM MgCl_2_), and (iv) neutral pH without Mg^2+^ (pH 7.0, 0 mM MgCl_2_).

Our SAXS measurements (**Figure 8A**) show that reducing pH has very little effect on the global conformation of 7SK SL-1, whereas the addition of Mg^2+^ drives 7SK SL-1 towards a more compact state (**Figure 8B-D**) – in excellent agreement with our simulated models, and previous SAXS measurements (19). In the absence of Mg^2+^, and regardless of pH, the Guinier *R_G_* sits between 25-26 Å (**Figure 8B**), in line with the extended NMR structure and our free energy surfaces. The addition of Mg^2+^ shifts the *R_G_* lower, to 21-22 Å (**Figure 8B**), which matches the value for the compact structures. The pair-distance distribution plots show the same trend, and provide a measure of the overall length of the RNA: addition of Mg^2+^ drives 7SK SL-1 into a shorter conformation regardless of pH, indicated by the lower *D_max_* value, and the more narrow *P(r)* curves (**Figure 8C**). In the presence of Mg^2+^ our SAXS measurements show 7SK SL-1 adopts a compact conformation with a *D_max_* of 76-78 Å. These measured values agree very well with the end-to- end distance of ∼76 Å observed in the crystallographic structures of the RNA (PDB IDs 5LYU, 5LYS) (17). In the absence of Mg^2+^, the *D_max_* is increased to ∼91-98 Å, consistent with the measured end-to-end distance of the elongated state solved by NMR (PDB ID 5IEM) (18).

**Figure 8.**
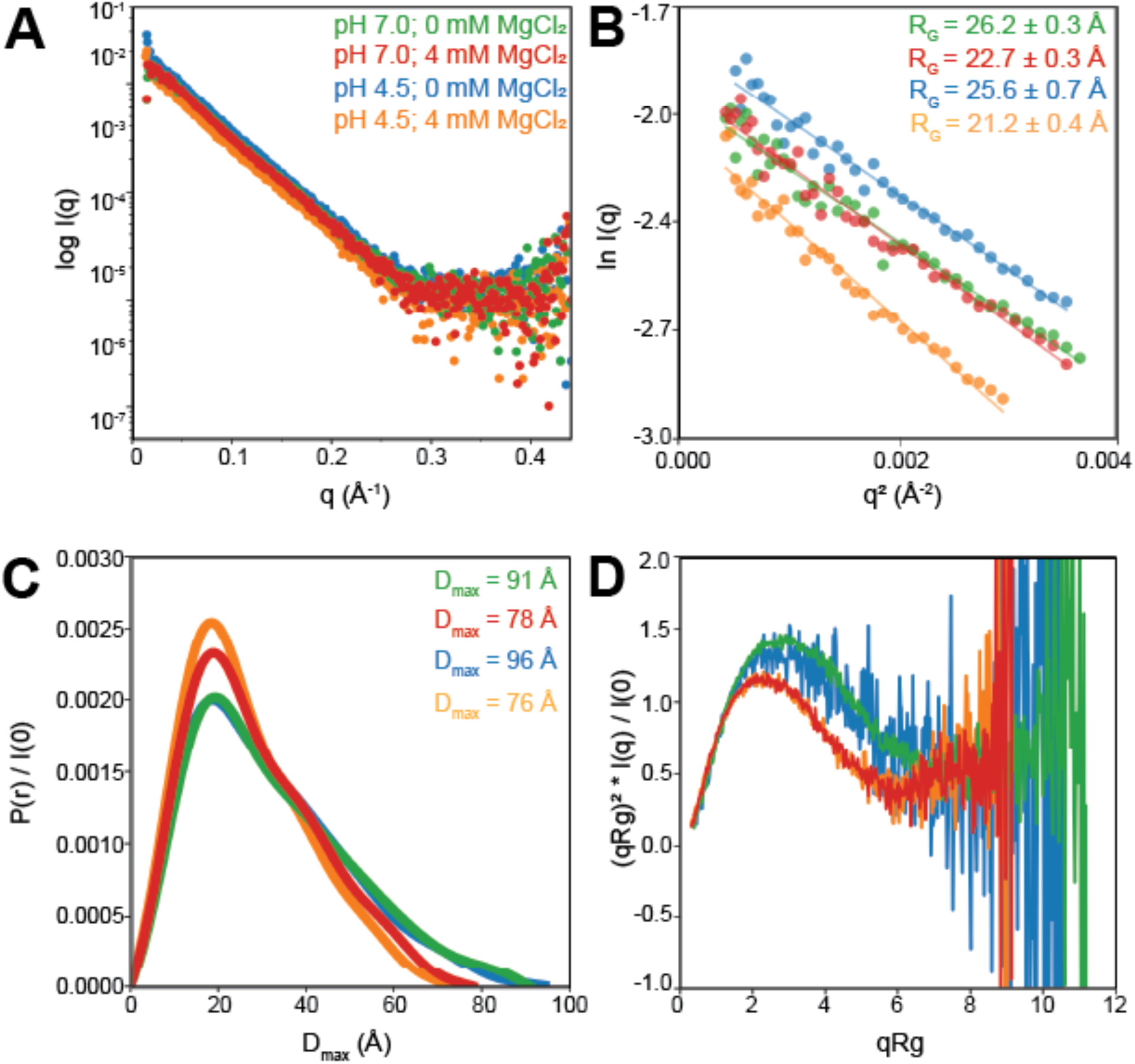
SAXS experiments deconvoluting the effect of pH and Mg^2+^ on the global conformation of 7SK SL-1. **A** SAXS scattering data, with calculated R_G_ values inlaid on the bottom left, **B** Guinier plots showing the fit-region and radius of gyration, **C** distance distribution plots, with D_max_ values indicated in the upper right, and **D** Kratky plots. All measurements were done on 7SK SL-1 at four conditions: pH 7.0, 0 mM MgCl_2_ (green), pH 7.0, 4 mM MgCl_2_ (red), pH 4.5, 0 mM MgCl_2_ (blue), and pH 4.5, 4 mM MgCl_2_ (orange).

To assess the structural integrity and flexibility of the 7SK SL-1 RNA constructs, we utilized a dimensionless Kratky analysis, plotting (qRg)^2^ I(q) / I(0) against qRg. Because the axes of this plot are normalized, they remain independent of molecular weight and overall size, allowing for a direct comparison of conformational states across different samples. For the RNA samples prepared in the presence of Mg^2+^ (both at pH 7.0 and pH 4.5), the profiles exhibited the classic signature of a well-folded biomolecule, characterized by a distinct local maximum near qRg ≈ √3, with a peak intensity of approximately 1.10. In contrast, the two Mg^2+^-free samples showed a clear deviation from this ideal folded behavior, with the (qRg)^2^ I(q) / I(0) values trending significantly higher (∼1.4) and the qRg values greater than the Mg^2+^-containing samples. These elevated values indicate a loss of the compact, globular shape seen in the folded state, suggesting that in the absence of magnesium, the RNA adopts a state of increased conformational flexibility and structural expansion (**Figure 8D**).

### NMR Experiments Show Local Structural Rearrangements

To monitor base triple formation, we carried out 2D ^1^H-^1^H nuclear Overhauser effect spectroscopy (NOESY) and 2D ^1^H-^15^N heteronuclear multiple quantum coherence (HMQC) experiments on two site-specifically labeled 7SK SL-1 constructs under varying pH and Mg^2+^ conditions. We focused on the U63 - G42:C67 and C71 - G74:C35 base triples (**Figure 9A**). Purines were ^15^N-labelled at the N1 position, and pyrimidines at the N1 and N3 positions. The imino proton fingerprint of the 7SK SL-1 constructs at pH 7.0 and 0 mM MgCl_2_ agrees well with previous NMR experiments, indicating proper folding of the RNA (**Supplemental Figure 9**) (18, 19). Chemical shifts and line broadening effects are observed around residues that cluster around the base triples in the 7SK SL-1 RNA.

**Figure 9.**
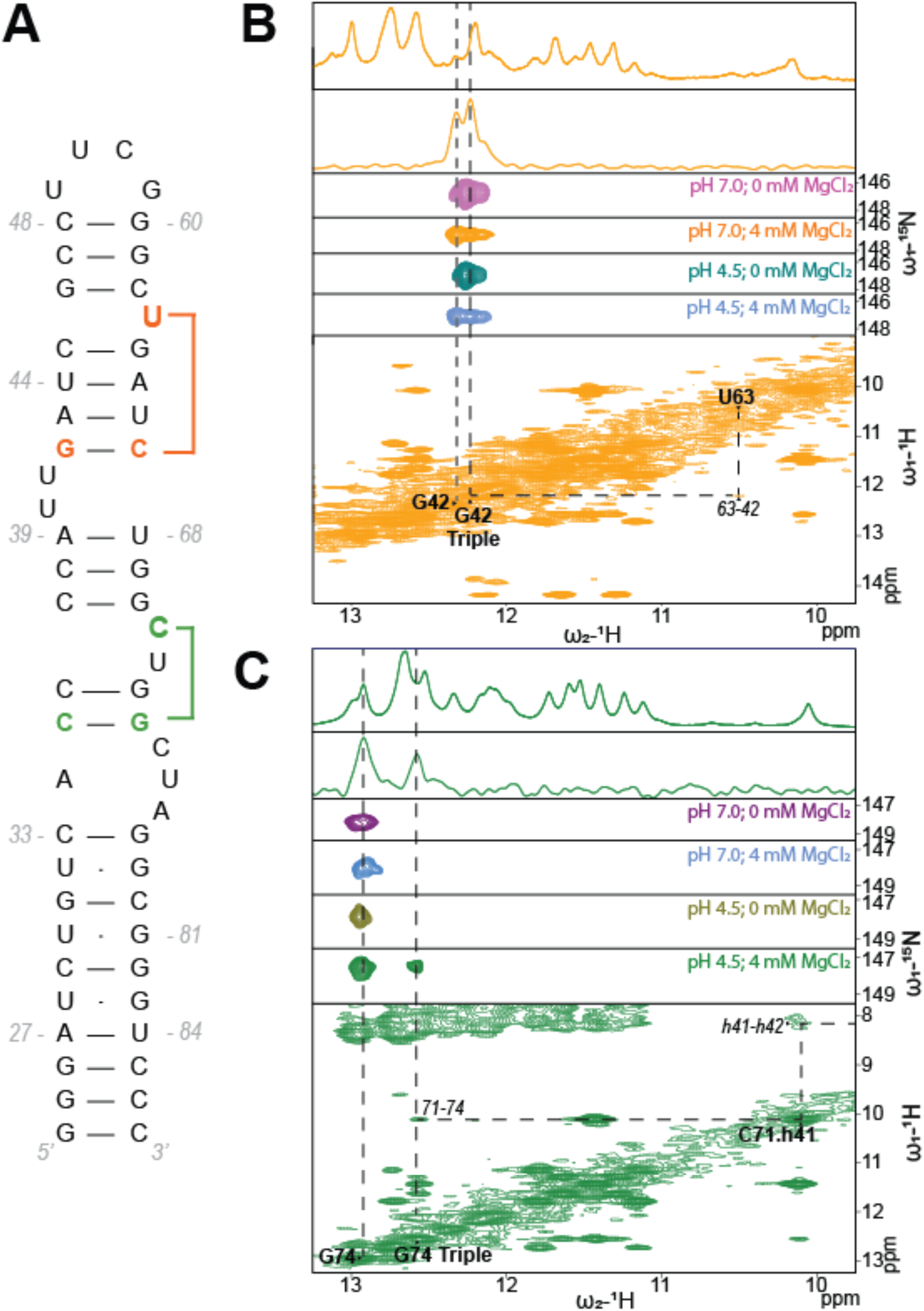
Measuring base triple formation using ^15^N selectively labeled constructs. **A** Secondary structure of the 7SK Sl-1 construct, indicating the selectively-labeled nucleotides involved in base triple formation. **B** 1D imino spectra, edited 1D ^1^H,^15^N-HMQC, 2D ^1^H,^15^N HMQC, and ^1^H,^1^H-NOESY spectra showing alignment between the U63-G42 NOE crosspeak, and the HMQC resonance belonging to the G42 triplex. **C** 1D imino spectra, edited 1D ^1^H,^15^N-HMQC, 2D ^1^H,^15^N HMQC, and ^1^H,^1^H-NOESY spectra showing an NOE crosspeak between the downshifted C71 amino, and the HMQC resonance belonging to the G74 triplex.

We first focused on the non-ionizable U63 - G42:C67 base triple. In the absence of Mg^2+^, both at pH 7.0 and pH 4.5, a shouldered peak corresponding to G42 is centered around 12.25 ppm (**Figure 9B**). We hypothesize that the main peak corresponds to the Watson-Crick-Francis G42:C67 base pair, and the shoulder reflects the sampling of the U63 - G42:C67 base triple (albeit at a lower population).

Upon addition of 4 mM MgCl_2,_ (pH 7.0 and 4.5), the G42 resonance undergoes a distinct chemical shift perturbation (**Figure 9B**). Specifically, the upfield peak, which corresponds to the base triple state, intensifies to match the downfield base-paired peak. This shift toward equal intensity suggests that Mg^2+^ directly stabilizes the triplex conformation, substantially increasing its population at equilibrium. This is further seen in the 1D imino-region, where addition of Mg^2+^ results in the formation of a new peak corresponding to the G42 triple (**Supplemental Figure 9**). Furthermore, we see several distinct CSPs upon the addition of Mg^2+^ (**Supplemental Figure 9C**), including G69, G70, G73, and G74 – the same series of sequentially stacked guanines as observed in our MD simulations, in excellent agreement with previous NMR studies reporting on CSPs resulting from the addition of Mg^2+^ to 7SK SL-1 (19).

Further corroborating these findings, we observe an NOE crosspeak between G42 and U63 in the presence of Mg^2+^ (**Figure 9B**), consistent with U63 - G42:C67 triple formation. Notably, this NOE peak is observed at both 2 and 4 mM MgCl_2_ (**Supplemental Figure 10**). The NOE between G42 and U63 is weak, likely due to (i) the dynamic nature of the base triple, and (ii) the G42:C67 pair residing at the edge of a bulge, where local fraying accelerates solvent exchange and induces broadening of the imino resonance. This conformational flexibility is reflected in our free energy surfaces, where the U63 - G42:C67 triple is sparsely populated, indicating a low thermodynamic barrier for escaping the triplex state (**Supplemental Figure 4**).

We next examined the behavior of the ionizable C71 - G74:C35 base triple, and found it is stabilized by low pH and Mg^2+^. At neutral pH, both in the presence and absence of Mg^2+^, a single G74 peak is observed in the HMQC spectra. At pH 4.5, in the absence of Mg^2+^, this single peak is also observed (**Figure 9C**). However, upon addition of Mg^2+^ at low pH, a second upfield resonance, corresponding to the base triple, is observed (**Figure 9C**). The presence of two distinct peaks indicates the triplex resides in a slow-to-intermediate exchange regime, sampling both base paired and base triple conformations on NMR timescales – consistent with our free energy surfaces showing 7SK SL-1 can spontaneously shift in and out of the triplex state with a minimal energetic penalty of 2-5 kcal/mol (**Figure 6**). Further supporting the formation of the base triple are the NOE connectivities between the H1 G74 base triple imino proton, to the severely downfield shifted N4 amino protons of C71 (**Figure 9C**). This downfield shift is characteristic of N3 protonation and formation of the triplex state (12, 16, 19).

Collectively, our NMR experiments demonstrate that both pH and Mg^2+^ remodel the free energy landscape of 7SK SL-1, shifting the conformational equilibrium to increase the population of the transiently sampled triplex states, in strong agreement with our calculated free energy surfaces.

## DISCUSSION

### Mg^2+^ and pH Have Distinct Effects on the Conformational Landscape of 7SK SL-1

We used MD simulations, free energy calculations, SAXS, and NMR experiments to probe the mechanism of base triple formation in 7SK SL-1, and deconvolute the effects of Mg^2+^ and pH on its conformational landscape.

Our MD simulations and free energy calculations rationalize how Mg^2+^ drives 7SK SL-1 towards more compact conformations. In our MD simulations, the Mg^2+^ ions interact with the phosphate backbone of 7SK SL-1 through their hydration spheres. The high-occupancy Mg^2+^ ions stabilize the compact state of 7SK SL-1 in two ways: either (i) by bridging the RNA backbone across the major groove through their hydration spheres, encouraging contraction of the RNA, or (ii) by stabilizing kinks in the RNA backbone near the base triples and crimping the backbone together (**Figure 3,4**). These interactions rigidify the RNA backbone of the bulged nucleotides, constraining them in a position more amenable for base triple formation, and help compensate for the steric strain resulting from the kinks in the RNA backbone needed to form the base triples. These results help explain observations from SAXS experiments, which show a compaction of 7SK SL-1 resulting from increased Mg^2+^ concentration, and more disordered behavior in the absence of Mg^2+^ (19). Despite the high-occupancy of these ion binding sites, the Mg^2+^ ions at these positions exchange rapidly in our simulations, and interact with the RNA primarily through their hydration spheres, which may explain why they were not observed in the X-ray structures.

At the local level, our free energy surfaces suggest that the non-ionizable base triples are stabilized only by Mg^2+^, whereas the ionizable base triples require both Mg^2+^ and protonation of the third base (**Figure 6**). This is validated by our NMR experiments, which show formation of the U63 - G42:C67 base triple requires *only* the presence of Mg^2+^ (**Figure 9A**). The C71 - G74:C35 base triple only forms at both lowered pH *and* in the presence of Mg^2+^ (**Figure 9B**), requiring protonation at the N3 position to maintain stable hydrogen bonding.

Our TI calculations estimate the *pKa* of the third nucleotides as high as ∼7.0 for C71 and C75, and up to ∼6.0 for A34 and A77. Based on these results, and available high-resolution structures, we hypothesized that only C71 and C75 are protonated at the near-neutral pH of the nucleus, and repeated our free energy surfaces with only C71 and C75 protonated. The free energy surface of base triple formation under these conditions shows the binding-competent, compact NMR structure as the most populated conformation compared to the base triple configuration observed in the X-ray structures (**Supplemental Figure 5**). Protonation of A34 and A77 in the X-ray structures may be short-lived excited states, or potentially an artifact of crystallization.

All of the base triples are stabilized by Mg^2+^, but only the four ionizable base triples are sensitive to pH. Both the ionizable and non-ionizable base triples require Mg^2+^ ions to form at high enough populations to observe through NMR. Conversely, lowered pH – that is, protonation of C71, C75, A34, or A77 – strongly increases the populations of the ionizable base triples, but has little effect on the non-ionizable base triples. Further, when C71 and C75 – and not A34 and A77 – are protonated in the presence of Mg^2+^, the binding-competent base triple configuration (PDB ID 6MCI) is more populated than the base triple configurations seen in the X-ray structures.

### Base Triple Formation Is Permitted Only From Specific Backbone Configurations

The third base of each base triple sits in a bulged region (**Figure 1**), each of which exhibits a kink in the RNA backbone when participating in a base triple. This kink compacts the RNA backbone and positions the third base within hydrogen bonding distance to its participating base pair. We quantified the change in the RNA backbone associated with compaction and base triple formation using the pseudo-torsion angles ***η*** and ***Θ*** (**Figure 7C, D**). Each base triple showed a narrowed ***η* / *Θ*** distribution upon formation (**Figure 7**; **Supplemental Figures 7-9**). This analysis demonstrates that base triple formation is only accessible from certain backbone conformations, involving local rearrangements in the RNA backbone across three nucleotides.

### A Proposed Mechanism for Base Triple Formation

Our results establish a foundation for understanding how the base triples of 7SK SL-1 form. The non-ionizable base triples appear to form through two sequential steps: (i) compaction of the RNA backbone at the bulged nucleotides proximal to the third base, and (ii) hydrogen bonding of the third base to the base triple. Because the base triples are only accessible from certain backbone conformations, rearrangement of the RNA backbone into a permissible conformation must occur before the third base can form a hydrogen bond to its participating base pair. The presence of Mg^2+^ facilitates this by stabilizing the electrostatic repulsion between phosphate groups in the kinked RNA backbone, restricting the third base in a conformational space amenable for base triple formation.

Formation of the ionizable base triples in 7SK SL-1 proceeds through three steps, the order of which is less clear: (i) compaction of the RNA backbone at the bulged nucleotides proximal to the third base, (ii) protonation of the third base, and (iii) hydrogen bonding of the third base to the base triple. It is ambiguous whether protonation of the third base occurs before or after compaction of the RNA backbone. Proton-coupled conformational changes have been previously reported in RNA (38). Future work will further examine the mechanism of proton-coupled base triple formation in 7SK SL-1, and dissect the rate-limiting step and overall kinetics of base triple formation in 7SK SL-1.

In both pathways, it is assumed that the participating base pair has already formed. However, if the participating base pair sits near a bulge, as observed with G78:C33 and G74:C35, the rate of base pair fraying potentially contributes to base triple formation.

### The Conformational Dynamics of 7SK SL-1 May Regulate its Binding with HEXIM

The relative populations of the different base triple configurations sampled by 7SK SL-1 may influence its binding with HEXIM and P-TEFb. Previous isothermal calorimetry experiments showed that Tat / HEXIM binding changes with Mg^2+^ and pH: increasing the amount of Mg^2+^ decreased non-specific binding for both Tat and HEXIM, whereas reducing the pH increased the binding affinity (19). We propose that the salt- and pH-dependence on binding is due to increased sampling of the base triples, which elevates the number of binding-competent conformations present in solution. Our free energy surfaces suggest that when C71, C75, A34, and A77 are protonated in the presence of Mg^2+^, 7SK SL-1 can spontaneously alternate between the different base triple configurations. The proton-coupled conformational flexibility of 7SK SL-1 could be a mechanism by which it regulates the transcriptional output of Pol II, by sampling both binding-competent and binding-incompetent conformations at an evolutionarily optimized rate dependent on cellular pH and Mg^2+^ concentrations.

Altogether, we demonstrate how pH and Mg^2+^ modulate the conformational dynamics of 7SK SL-1, and provide a mechanism for base triple formation at atomic detail. This work expands our textbook knowledge on RNA structure and dynamics, and establishes a framework for probing base triple dynamics in RNA. As the number of high-resolution RNA structures increases, it will be interesting to see if other RNAs harboring base triples exhibit similar dependencies on pH and Mg^2+^ in the formation of base triples.

## METHODS

### Simulation Protocols

Simulations of 7SK SL-1 were carried out using the PMEMD module of the Amber v23 software package on v100 graphical processing units (GPUs). The *LJbb* parameter set was used to model the RNA; this force field combines the *OL3* force field with modified phosphate oxygen van der Waals parameters from Steinbrecher and Case (39, 40). The *OPC* water model was used to describe the solvent, and the Li–Merz 12-6-4 ion parameters were used for the mono-and divalent ions (41, 42). The long-range Coulombic interactions were calculated using the particle-mesh Ewald method with a 10.0 Å cutoff (43). Bonds involving hydrogen atoms were restrained using the SHAKE algorithm with a 0.00001 Å tolerance (44). All production simulations were carried out in the NPT ensemble using a Berendson barostat at 1.0 atm, with Langevin dynamics using a 1.0 ps^-1^ collision frequency, a 4.0 fs timestep, and hydrogen mass repartitioning (45–47). Structure visualization was conducted using VMD, and analysis was carried out using the *cpptraj* module of AmberTools v23 (48–50).

### Molecular Dynamics Simulations of 7SK SL-1

MD simulations of 7SK SL-1 were performed starting from the each of the four experimental structures: the crystal structure in the IN conformation (PDB ID 5LYU; Assembly 1; model A), the crystal structure in the OUT conformation (PDB ID 5LYS; Assembly 1; model B); the NMR structure in the compact state (PDB ID 6MCI; model 1); and the NMR structure in the extended state (PDB ID 5IEM; model 1) (16–18).

Hydrogen atoms were added if missing from structures and the systems were solvated in a 16 Å truncated octahedron periodic box with neutralizing K^+^ counter-ions and a 5 mM MgCl_2_ buffer using the *tleap* module of AmberTools 2023 (50). The final system size was 72,333 atoms. Each system was simulated at low and neutral pH, both with and without Mg^2+^. The “low-pH” systems described in this text refer to systems where A34, C71, C75, and A77 are protonated. After building, the systems were equilibrated using a 10-step equilibration protocol.

The water molecules were first minimized for 1,000 steps using steepest descent and then for an additional 9,000 steps with conjugate gradient, while the rest of the system was restrained with 1 kcal/(mol*Å^2^) Cartesian positional restraints. The systems were then heated to 298.15 K at constant volume over 0.5 ns using 100 kcal/(mol*Å^2^) positional restraints applied to all atoms except hydrogens and water molecules. The box size and density were equilibrated over 1 ns using a constant pressure of 1 bar with the same positional restraints as the previous step. The restraints were then lowered to 10 kcal/(mol*Å^2^) for an additional 1 ns of equilibration. A second minimization step was carried out using 10,000 steps of conjugate gradient minimization with atoms C3’, C4’, C5’, O5’, P, and O3’ restrained with a 10 kcal/(mol*Å^2^) force constant. The next five steps were done at constant NPT for 20 ns each with positional restraints on the same atoms, reducing the force constant at each step from 10 kcal/(mol*Å^2^), to 5, to 2.5, to 1.0, to 0.5 kcal/(mol*Å^2^), respectively. Equilibration simulations were done using a 2 fs timestep. Production simulations were carried out in triplicate for each of the three systems for 2.5 µs with a 4 fs time step and hydrogen mass repartitioning; the last 2.0 µs was used for analysis (47). Four structures simulated in four conditions, in triplicate, with 2.0 µs of production simulation giving 96.0 µs of MD simulation time.

### pKa Calculations Using Thermodynamic Integration

The TI calculations were performed in one step at 298 K using a soft-core potential as implemented in Amber23 by disappearing the proton in C^+^ / A^+^ to the neutral form of each nucleotide, starting from each experimental structure. The transmutation was done either in the context of the RNA structure, or as a free methyl-capped nucleotide in a box of water (see **Supplemental Figure 3** for the free energy cycle used). A 1 fs time step was used for the TI calculations. The TI calculations were carried out with a series of λ-values = 0.00922, 0.04794, 0.11505, 0.20634, 0.31608, 0.43738, 0.56262, 0.68392, 0.79366 0.88495, 0.95206, and 0.99078. Each λ window was simulated for 100 ns using the CUDA variant of PMEMD (50). TI calculations were done twice with different initial ion positions. The pKa values were calculated by averaging the first and second 50 ns of the two 100 ns calculations. The uncertainty is the standard deviation of the four 50 ns segments. The trapezoidal rule was used for the integration of all λ windows.

### Well-Tempered Adaptively Biased Molecular Dynamics Simulations

To calculate the free energy surface of base triple formation with and without the third bases protonated, we used the WT-ABMD algorithm as implemented in Amber v23 (34). The WT-ABMD algorithm has been applied previously to calculate free energy surfaces of conformational changes of both protein and nucleic acid systems (51–55). The ABMD method is a form of umbrella sampling that uses a history-dependent biasing potential, and thus requires the usage of CVs to calculate the free energy surface. We applied WT-ABMD using the Radius of Gyration (*R_G_*) and the number of base triples as the two CVs. The atomic coordinates of all non-hydrogen RNA atoms were used as the input mask for the *R_G_* CV.

The number of base triples was measured using the coordination number CV, which is defined in Amber as:

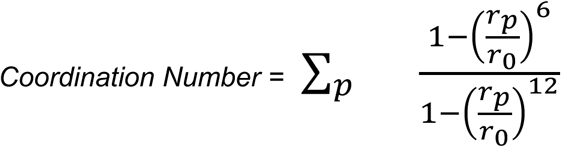

where *r_p_* is the distance between atoms of pair *p*, and *r_0_* is the defined reference distance in Å. We used the distance between the N3 atoms of pyrimidines and N7 atoms of purines in each base triple as the pairs of atoms for the coordination number CV. The *r_0_* reference distance was defined as 3.5 Å. The exact atom pairs used are summarized in **Table 1**. Note that, even if the *r_p_* value is less than the reference value *r_0_* an integer value of the CN is not ensured, explaining the non-integer values of CN seen in the free energy surfaces.

**Table 1.**
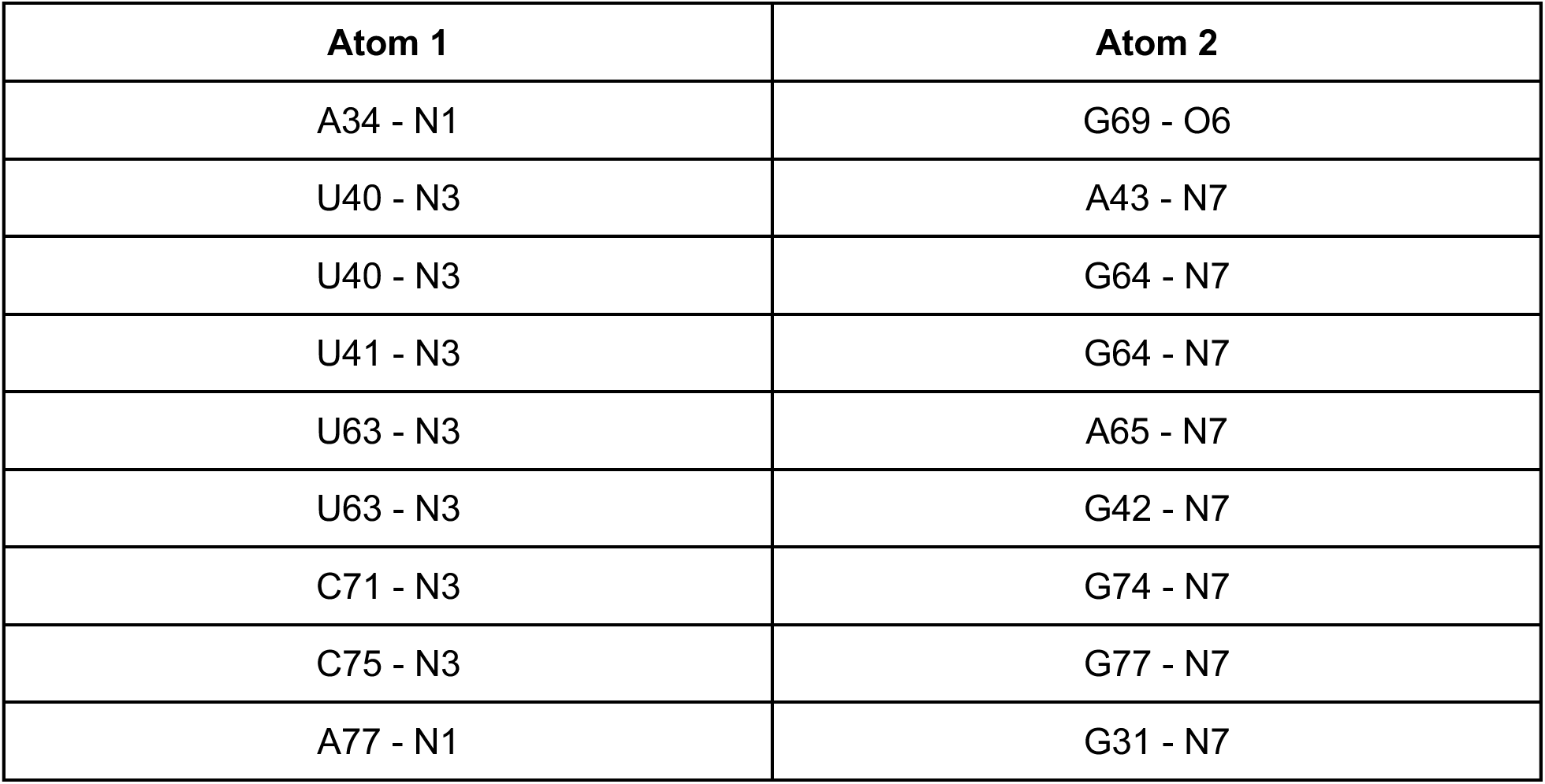
The atom pairs used in the coordination number CV for WT-ABMD, used to measure the number of base triples.

We used the WT-ABMD method with 8 replicas at 298 K. The final structures immediately following the equilibration protocol were used as the starting structures for the WT-ABMD simulations, with two replicas per structure. The pseudo-temperature, resolution, and *τ*_F_ were set to 10,000 K, 0.08 Å, and 10.0 ps, respectively. The WT-ABMD simulations were done in 50 ns “cycles,” where each cycle was repeated until the resulting PMFs no longer changed after addition of more cycles. We conducted 25 cycles of WT-ABMD for both the protonated and deprotonated systems, totaling 10.0 µs of simulation time for each free energy surface. We repeated the WT-ABMD simulations with only C71 and C75 protonated using the same protocol. The PMFs were calculated using the *nfe-umbrella-slice* utility of AmberTools v23.

We ensured convergence of our free energy surfaces by conducting multiple cycles of WT-ABMD (8 parallel replicas, 25 cycles, 50 ns per cycle – for a total of 10.0 µs per free energy surface), repeating until the resulting PMF no longer changes.

### RNA Samples Preparation

Unlabelled RNA for SAXS and NOESY experiments was prepared by in vitro transcription using in-house T7 RNA polymerase from oligonucleotide templates purchased from IDT. The RNA was purified on 8M urea denaturing acrylamide gels followed by electroelution, desalting, and lyophilization. Samples were resuspended in the chosen buffer for NMR or SAXS measurements, and refolded by heating at 95°C and snap-cooled on ice. Concentrations were measured using a Nanodrop Spectrometer.

### SAXS Measurements

SAXS data were collected on a Xenocs BioXolver at Cornell University. Four samples at concentrations of 0.5, 1.0, and 4.0 mg/mL were prepared in 15 mM NaPi and 25 mM NaCl, and one of four buffer conditions: (i) pH 4.5, 0 mM MgCl_2_; (ii) pH 4.5, 4 mM MgCl_2_; (iii) pH 7.0, 0 mM MgCl_2_; (iv) pH 7.0, 4 mM MgCl_2_. Buffer subtraction, scaling, and merging of data was performed using BioXTAS Raw (56).

### NMR Spectroscopy

All NMR experiments were carried out using 250 uM ^15^N selectively-labeled RNA for ^1^H 1D imino experiments with standard Bruker pulse programs on a Bruker 800 MHz Avance III spectrometer equipped with a TCI cryogenic probe. The RNA was dissolved in a buffer containing 15 mM sodium phosphate and 25 mM NaCl. Several samples were prepared at pH values of 4.5 and 7.0, with either 0, 2, or 4 mM MgCl_2_. Spectra for observing exchangeable protons were conducted at 283 K in 10% D_2_O. For NOESY experiments, the mixing time was set to 200 ms. ^1^H spectra were processed and analyzed using Bruker TopSpin (4.4) and CCPN (3.4.1) (57).

## DATA AVAILABILITY

All study data are included in the article and/or supporting information. SAXS data have been deposited in the SASBDB under accession codes SASDZE6, SASDZF6, SASDZG6, SASDZH6.

## ACKNOWLEDGEMENTS

Support for the work performed at Cornell University was provided by the Pollack lab at Cornell University. The authors thank Sarah Uttormark, Qingyue Hu, and Hugh Higinbotham for experimental assistance, and Dr. Lois Pollack for access to instrumentation and insight during manuscript preparation. Selectively-labeled RNA samples were provided by Raphael Plangger from the Kreutz Lab at the University of Innsbruck. Computational research was conducted on the Torch High Performance Computer at New York University (NYU), supported by the NYU High Performance Computing resources, services, and staff expertise. NMR spectroscopy experiments were carried out at the NYU Shared Instrumentation Facility, with support from Dr. Chin Lin and Dr. Joel Tang. The authors further thank Dr. David Klingler for assistance with NMR experiments.

## AUTHOR CONTRIBUTIONS

L.F. Investigation, conceptualization, visualization, data analysis, writing original draft.

H.V. Investigation, writing original draft.

A.N.J. Conceptualization, writing original draft, funding acquisition, resources, supervision.

