## Supplemental Information for "A Shifting Landscape: Deconvoluting the Effect of Mg^2+^ and pH on Base Triple Formation in 7SK SL-1"

**Supplemental Note 1 -** Radius of gyration (*R_G_*) measures the global compactness of a biomolecule, with larger values corresponding to more extended states, and lower values corresponding to more compact states.

**Supplemental Note 2** - The Adaptively Biased Molecular Dynamics (ABMD) method is a mature simulation algorithm that has been applied to protein and nucleic acid systems for the determination of free energy surfaces (37–40). ABMD is a form of umbrella sampling that uses a history-dependent biasing potential, and requires the use of collective variables (CVs) (41).

**Supplemental Figure 1.** Distance distributions of the base triples throughout triplicate 2.0 µs MD simulation of 7SK SL-1 starting from each of the four experimental structures, with Mg^2+^ ions, at both low (grey) and neutral (colorful) pH values. The base triples are colored to match the color scheme from Figure 1. The MD simulations were initialized starting from the compact solution NMR structure (PDB ID 6MCI), the compact X-ray structures (PDB ID 5LYU and 5LYS, respectively), and the extended solution NMR structure (PDB ID 5IEM). The blue dots indicate the native base triples in each experimental structure.

**Supplemental Figure 2.** Distance distributions of the base triples throughout triplicate 2.0 µs MD simulation of 7SK SL-1 starting from each of the four experimental structures, with no Mg^2+^ ions, at both low (grey) and neutral (colorful) pH values. The base triples are colored to match the color scheme from Figure 1. The MD simulations were initialized starting from the compact solution NMR structure (PDB ID 6MCI), the compact X-ray structures (PDB ID 5LYU and 5LYS, respectively), and the extended solution NMR structure (PDB ID 5IEM). The blue dots indicate the native base triples in each experimental structure.

**Supplemental Figure 3.** Free energy cycle constructed for the calculation of the pKa shift for the third base in the ionizable base triples.

**Supplemental Figure 4.** Distance distributions and R_G_ of each base triple in the WT-ABMD simulations. **A** Low pH with Mg^2+^, **B** neutral pH with Mg^2+^, **C** low pH without Mg^2+^, and **D** neutral pH without Mg^2+^.

**Supplemental Figure 5.** PMF of 7SK SL-1, with *only* C71 and C75 protonated, alternating between the four different experimental structures with Radius of Gyration and Coordination Number as the two CVs. The black stars indicate the values of the experimental reference structures, and their PDB IDs are labeled for clarity. Each color change represents a 2.5 kcal/mol change in free energy, while contour lines represent a 5.0 kcal/mol change in energy.

**Supplemental Figure 6.** Eta / theta (*η* / *Θ)* distributions of C75, U76, and A77 for structures from our ABMD simulations in which the C75 base triple is **A** present, or **B** absent. The population is colored according to intensity.

**Supplemental Figure 7.** Eta / theta (*η* / *Θ)* distributions of A34, C35, and C36 for structures from our ABMD simulations in which the A34 base triple is **A** present, or **B** absent. The population is colored according to intensity.

**Supplemental Figure 8.** Eta / theta (*η* / *Θ)* distributions of U76, A77, and G78 for structures from our ABMD simulations in which the A77 base triple is **A** present, or **B** absent. The population is colored according to intensity.

**Supplemental Figure 9.** Imino spectra of 7SK SL-1 under different conditions. **A** 1D imino spectra of 7SK SL-1 at (red) pH 7.0 with 0 mM MgCl_2_, and (blue) pH 7.0 with 4 mM MgCl_2_. **B** 1D imino spectra of 7SK SL-1 at (green) pH 4.5 with 0 mM MgCl_2_, and (purple) pH 4.5 with 4 mM MgCl_2_. **C** Secondary structure of 7SK SL-1, with nucleotides that undergo CSPs upon addition of Mg^2+^ colored in red.

**Supplemental Figure 10**. The U63-G42 NOE crosspeak is visible at 2 mM MgCl_2_. ^1^H,^15^N HMQC, and ^1^H,^1^H-NOESY spectra showing alignment between the U63-G42 NOE crosspeak, and the HMQC resonance belonging to the G42 triplex at neutral pH and 2 mM MgCl_2_.
